# Plant genome response to incoming coding sequences: stochastic transcriptional activation independent of integration loci

**DOI:** 10.1101/2020.11.28.401992

**Authors:** Soichirou Satoh, Takayuki Hata, Naoto Takada, Makoto Tachikawa, Mitsuhiro Matsuo, Sergei Kushnir, Junichi Obokata

**Author notes:** These authors contributed equally to this work (Soichirou Satoh, Takayuki Hata). Soichirou Satoh and Takayuki Hata should be considered joint first author.

## Abstract

Horizontal gene transfer can occur between phylogenetically distant organisms, such as prokaryotes and eukaryotes. In these cases, how do the translocated genes acquire transcriptional competency in the alien eukaryotic genome? According to the conventional view, specific loci of the eukaryotic genome are thought to provide transcriptional competency to the incoming coding sequences. To examine this possibility, we randomly introduced the promoterless luciferase (LUC)-coding sequences into the genome of *Arabidopsis thaliana* cultured cells and performed a genome-wide “transgene location vs. expression” scan. We mapped 4,504 promoterless *LUC* inserts on the *A. thaliana* chromosomes, and found that about 30% of them were transcribed. Only a small portion of them were explained by the conventional transcriptional fusions with the annotated genes, and the remainder occurred in a quite different manner; (1) they occurred all over the chromosomal regions, (2) independently of the insertion sites relative to the annotated gene loci, inherent transcribed regions, or heterochromatic regions, and (3) with one magnitude lower transcriptional level than the conventional transcriptional fusions. This type of transcriptional activation occurred at about 30% of the inserts, raising a question as to what this 30% means. We tested two hypotheses: the activation occurred at 30% of the entire chromosomal regions, or stochastically at 30% of each insertion event. Our experimental analysis indicates that the latter model could explain this transcriptional activation, a new type of plant genome response to the incoming coding sequences. We discuss the possible mechanisms and evolutionary roles of this phenomenon in the plant genome.

## Introduction

Horizontal or endosymbiotic gene transfer (HGT/EGT) events greatly contributed to the evolution and diversification of terrestrial life and organisms [1, 2]. In the plant phyla, thousands of genes originally encoded by photosynthetic bacteria were transferred to the host nuclear genome to produce photosynthetic eukaryotes [3], even though the bacterial genome systems were quite different from those of eukaryotes. The genetic flow from the plastid to the nuclear genome is ongoing [4, 5].

However, the mechanisms by which foreign genes obtained by HGT/EGT events acquired transcriptional competency in the host nuclear genome remain largely unclear. Examination of the transferred DNA fragments from the plastid to the nucleus led to the proposal that the translocated genes were transcribed as fusion transcripts via trapping of endogenous genes/promoters [6, 7]. This transcriptional activation mechanism is easy to understand, but one promoter-acquisition event will result in one disruption of a preexisting gene. Therefore, it has a shortcoming with regard to explaining the thousands of functional gene transfers that occurred in the endosymbiotic evolution.

The cryptic promoter hypothesis could be an alternative to explain the transcriptional activation of exogenously incoming genes in the nucleus. This hypothesis was originally adopted to explain the enigmatic expression of the coding sequences in gene-/promoter-trapping experiments; promoterless coding sequences were occasionally transcribed without obvious trapping of any annotated genes/promoters [8–14]. The cryptic promoter hypothesis postulated that the invisible promoters hidden in the genome, namely cryptic promoters, capture the incoming coding sequences to cause their detectable transcription. However, molecular identities of these cryptic promoters have long been unsolved. Recently, we demonstrated that such unexpected transcriptional activation in gene-/promoter-trapping experiments occurred via at least two different mechanisms in the plant genome: (1) cryptic promoter capturing, in which exogenous DNA was transcribed by trapping a preexisting promoter-like chromatin configuration that is not associating with annotated genes; and (2) promoter *de novo* origination, in which promoter-like epigenetic landscapes were newly formed via chromatin remodeling triggered by the insertion of a coding sequence [15]. We should note that these two mechanisms could endow transcriptional activity to the incoming coding sequences without disturbing the preexisting nuclear gene network. In examining whether these cryptic promoters could be a source of transcriptional activation in massive gene transfer, we should know how often the cryptic promoter activation occurs in the whole nuclear genome.

In this study, we applied a massively parallel reporter assay [16, 17] to the conventional gene-/promoter-trapping experiments and carried out a genome-wide “transgene location vs. expression” scan. We introduced thousands of promoterless coding sequences of firefly luciferase (LUC) genes as a model of transferred genes into the genome of *Arabidopsis thaliana* cultured cells, and examined the manners by which transcriptionally inert transgenes become activated in the foreign genome environment. We found that a small portion of the transcriptional activation of transgenes was explained by the conventional gene-/promoter-trapping mechanism, but the majority of promoterless *LUC* inserts were transcriptionally activated in a quite different manner, i.e., integration-dependent stochastic transcriptional activation. This transcriptional activation occurred stochastically at about 30% of each insertion event, independently of the integration locus relative to the preexisting genes, inherent transcribed regions, or heterochromatic regions. We discuss the likely mechanism of this transgene activation phenomenon and refer to its possible contribution to the initial transcriptional activation process of HGT/EGT during plant genome evolution.

## Results

### General view of the transgene expression over the entire genome

To understand the rules that govern the transcriptional activation of alien incoming genes, we introduced thousands of promoterless luciferase (*LUC*) genes into *A. thaliana* T87 suspension-cultured cells via *Agrobacterium*-mediated transformation (S1A Fig). In the pools of transformed cells, 4,504 *LUC* genes were mapped onto the *A. thaliana* genome (Fig 1A, and S1B, C, and S2 Figs). The *LUC* genes were evenly distributed across the length of the five *A. thaliana* chromosomes, with the exception of the pericentromeric regions, where the insertion frequency was significantly lower (Fig 1A). The relative abundances of the *LUC* genes inserted in the intergenic, genic, and promoter regions were roughly proportional to the relative lengths of these genomic regions (Fig 1B and S3 Fig). On the fine distribution map of the inserts, genic promoter regions (~200 bp) were more prone to be inserted than the other regions by about threefold (Fig 1B, and S3 and S4 Figs), in accordance with a relatively open chromatin configuration of the promoter region. Despite such slight biases, the *LUC*-mapped loci covered entire chromosomal regions (Fig 1A and B), and thus were suitable for the genome-wide scanning of transgene transcriptional activation events.

**Fig 1.**
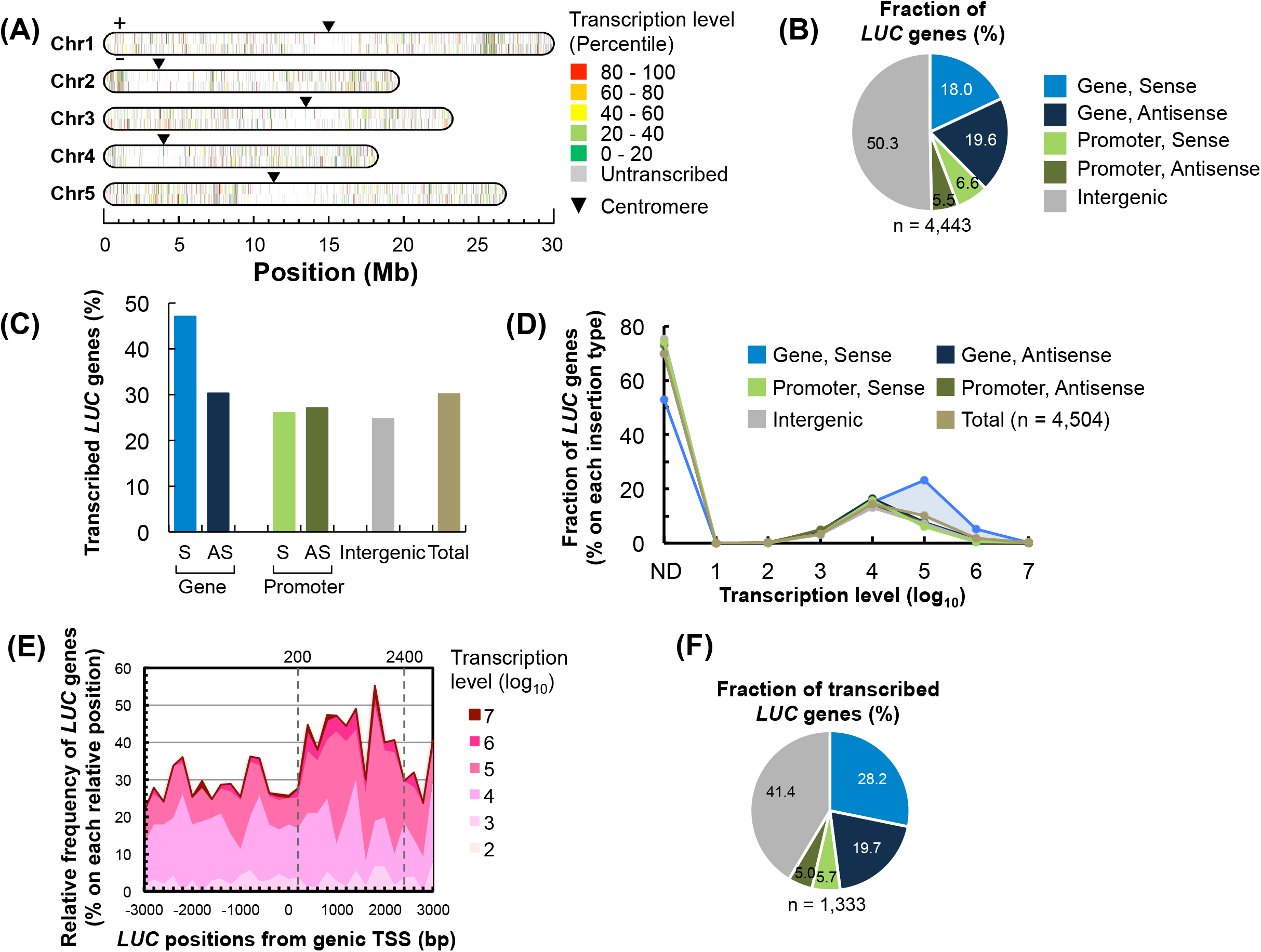
Massively parallel promoter-trapping analysis of the *Arabidopsis thaliana* genome. (A) Genomic positions of the inserted promoterless *LUC* genes and respective transcription levels. The bars represent the 4,504 mapped *LUC* genes regarding their orientation towards the upper (+) or bottom (–) DNA strands of the five *A. thaliana* chromosomes. The colour scheme discriminates *LUC* genes according to their expression levels. (B) Relative abundance of the *LUC* gene insertion types relative to the annotated gene locations. The *LUC* genes that cannot be classified their insertion type uniquely were omitted. (C) Percentage of transcribed *LUC* genes within the respective insertion types. S and AS indicate the sense and antisense orientations, respectively. (D) Distribution profiles of the *LUC* genes of respective insertion types according to the transcription level, with the total frequency of each insertion type normalized to be 100%. The light-blue area indicates the superposed fraction in the genic-sense insertion type. (E) Abundance of the *LUC* genes with the indicated transcription levels in relation to the distance from the genic TSS in each window (200 bp). (F) Classification of the transcribed *LUC* genes according to their insertion types, as in (B).

RNA deep sequencing was used to detect and measure the *LUC* expressions. Unique barcode identifiers enabled us to link the distinct genomic locations and *LUC* transcription levels at each position (Fig 1, and S1B and D Fig). We found that 1,355 of the 4,504 *LUC* genes identified were transcribed with a ~10^5^-fold variation in *LUC* mRNA levels (Fig 1D). Some barcodes could possibly behave as *cis*-regulatory elements and affect their own expression. However, our correlation analyses did not provide evidence of such function of barcode sequences (S5 and S6 Figs).

### Identification of two distinct mechanisms of transgene transcriptional activation

In the simplest-case scenario, promoterless *LUC* transcription is a result of the trapping of endogenous transcription units. To test this conventional model, we classified the 4,504 *LUC* loci into five insertion types in relation to the annotated genes: (i) sense and (ii) antisense orientation within the gene-coding regions, (iii) sense and (iv) antisense orientation in the promoter regions, and (v) intergenic regions. According to this classification, 25–30% of the *LUC* genes in each insertion type were transcribed, except for the genic-sense insertion type; about 50% of them were transcribed in the genic-sense fraction (Fig 1C). Why are the genic-sense inserts more prone to be transcribed?

As shown in Fig 1D, the transcription levels of *LUC* genes in each insertion type ranged from 10^1^ to 10^7^ at the mean transcription level of 10^4^, with that of the genic sense type exceptionally high, at the level of 10^5^. The comparison of the distribution profiles of the five insertion types revealed that the genic-sense type had a superposed fraction (light-blue fraction in Fig 1D) at higher transcription levels (10^5^–10^7^). Without this superposed fraction, the distribution curves of the five *LUC* insertion types were remarkably similar (Fig 1D). To explain this result, we next examined the *LUC* insertion sites relative to the annotated genic transcription start sites (TSSs) and *LUC* transcription levels (Fig 1E). We found that the *LUC* inserts with higher transcription levels (10^5^–10^7^) were more abundant at 0.2–2.4 kb downstream of the annotated TSSs (Fig 1E). Without this superposed fraction in this region, the expressed inserts appeared to be similarly distributed both within and outside of the annotated transcribed regions (S7 Fig). In *A. thaliana*, the median lengths of the 5’ untranslated regions (UTRs) and mRNAs are ~70 and ~1,900 bp, respectively (as calculated from the TAIR10 database, https://www.arabidopsis.org/index.jsp); thus, the region 0.2–2.4 kb downstream from the annotated TSS roughly corresponds to the intrinsic protein-coding regions. Based on these observations, the *LUC* inserts of the genic-sense type appeared to be transcribed at least in part by the conventional gene-trapping mechanism, in addition to the transgene transcription mechanism that similarly occurred over the entire genome.

If our above assumption is the case, the contribution of the conventional gene-trapping to the whole transcriptional activation of the incoming coding sequences is small; rather, the majority of transcriptional activation occurred by the distinct mechanism, even within the genic-sense insertion type (Fig 1D-F). The mean transcription level of this transcriptional activation was 10^4^, which was one magnitude lower than that of the conventional transcriptional fusions (Fig 1D and E). To confirm that this whole expression profile was not a sequencing artifact, we performed similar analyses using more reliable datasets (i.e., the *LUC* inserts whose sequencing reads were more highly abundant than the background level) with elevated read number threshold. Irrespective of the threshold read numbers, two distinct fractions corresponding to the gene-trapping type (light-blue fraction in S8A and B Fig) and the other type (light-red fraction in S8A and B Fig) were clearly detected, as in Fig 1D. In addition, these distribution profiles were confirmed by three biologically independent samples (S8C-E Fig). Based on these analyses, we concluded that the low-level transcriptional activation of transgenes that occurred over the entire chromosomal regions was not a sequencing artifact.

### Promoterless *LUC* genes were transcribed regardless of inherent transcriptional activities

Pervasive transcription throughout the genome characterizes eukaryotic organisms. We asked whether the genome-wide transcription of the *LUC* genes could be explained by the integration within such pervasively transcribed regions. To define the genomic transcription landscape of the *A. thaliana* T87 cells studied here, we performed deep RNA sequencing of the wild-type (WT) cells. We classified the 4,504 *LUC* loci by comparing their transcription status between transgenic and WT cells (Fig 2A). Unexpectedly, only 7.8% of the *LUC* genes were transcribed in the inherently transcribed genomic regions (type (iii) in Fig 2A), whereas 22.3% of the *LUC* genes were transcribed in the transcriptionally inert regions (type (i) in Fig 2A). As for the 7.8% of the *LUC* genes (type (iii) in Fig 2A), we compared the transcription levels between the transgenic and WT cells, but no correlation was found (Fig 2B, *r* = 0.21). Two conclusions were drawn from this analysis: (1) transcriptional activation of the *LUC* inserts occurs independently of the inherent transcriptional status of the genomic region where the *LUC* was inserted; and (2) the transcriptional activities of the *LUC* inserts do not reflect the inherent transcriptional activities of the given genomic regions.

**Fig 2.**
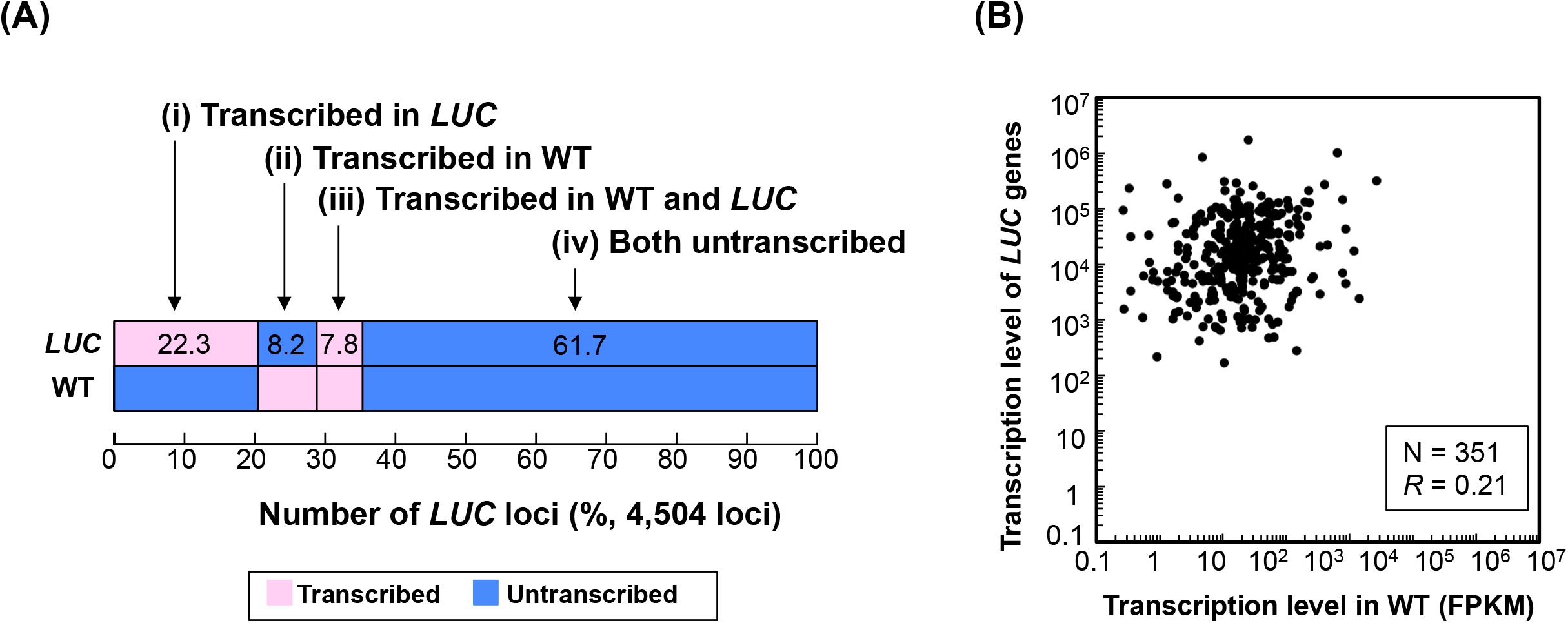
Transcription states of the *LUC* loci in WT and transgenic cells. (A) The 4,504 *LUC* loci were clustered into four groups according to the combination of on/off transcription states in WT and transgenic cells. The local transcription landscape in WT cells was determined based on the RNA-Seq analysis. (B) Comparison of the transcription levels between WT and transgenic cells for the *LUC* loci that were transcribed in both WT and transgenic cells.

### Transcriptional activation of promoterless *LUC* genes was not affected by the inherent heterochromatic status

We wondered whether *LUC* transgenes could overcome the silencing effects of the histone code. In *A. thaliana*, the dimethylation of the ninth lysine residue of histone H3 (H3K9me2) is thought to be associated with transcriptional silencing in the heterochromatic regions [18–20]. A Chromatin immunoprecipitation-sequencing (ChIP-Seq) analysis of the WT *A. thaliana* T87 cells revealed that 15.6% of the genome was covered by H3K9me2-containing chromatin and was largely associated with pericentromeric regions. In the transgenic cells, only 120 *LUC* genes were inserted into the H3K9me2-containing heterochromatic regions (see the legend of Fig 3), indicating that the integration frequency in this region was one-seventh of the rest of the genome. However, in the H3K9me2-containing region, 28% of the *LUC* inserts were transcriptionally activated and their activation profiles were similar to the other regions (Fig 3A). The transcription levels of these *LUC* genes did not show any correlation with the degree of H3K9me2 modification (Fig 3B). Furthermore, two transcribed *LUC* genes were located 63 kb and 682 kb from the centromeres (S9 Fig), and these regions were covered by pericentromeric heterochromatin [21]. Taken together, we concluded that the transcriptional activation of the *LUC* inserts occurred at a rate of about 30% irrespective of the inherent heterochromatic status.

**Fig 3.**
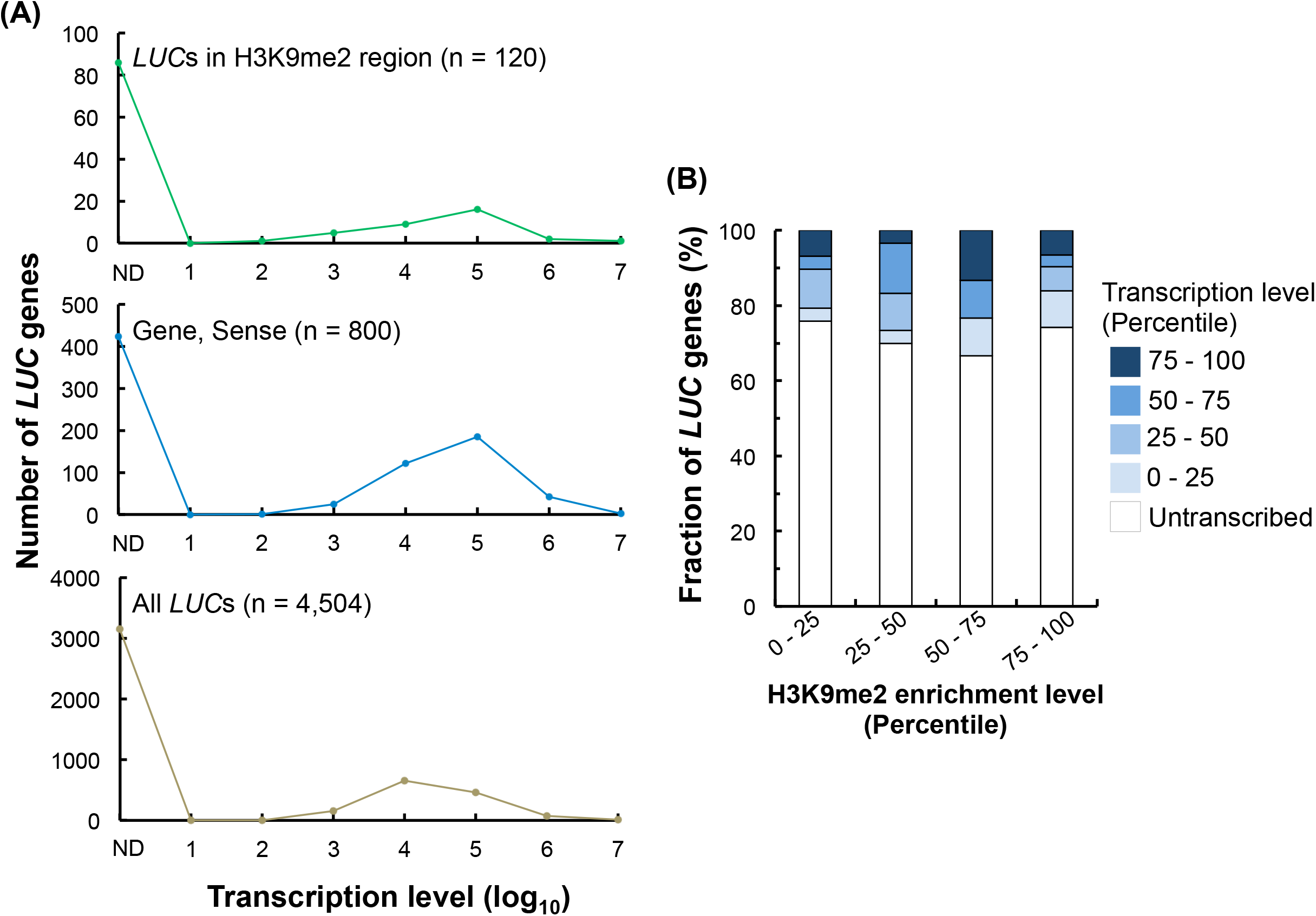
Transcription states of the *LUC* genes in the heterochromatic regions. (A) The upper panel shows the transcription profile of the *LUC* genes in the heterochromatic regions. The middle and bottom panels are derived from Fig 1D and represent the transcription profiles of the genic-sense type and all of the *LUC* genes, respectively. H3K9me2-marked heterochromatic regions covered 18.6 Mb in total and accounted for ~15.6% of the genome, where 120 *LUC* genes were inserted. About 80% of the H3K9me2-marked regions lay within the pericentromere. (B) Transcription levels of the *LUC* genes relative to the increased enrichment of H3K9me2. The transcription levels and H3K9me2 enrichment are both shown as percentiles based on all of the *LUC* genes located in the H3K9me2-marked heterochromatic regions.

### Integration-dependent stochastic activation of transgene transcription

As described above, transgene transcriptional activation was observed for 30% of the *LUC* inserts, which raised a question: What does this 30% mean? To account for this question, we hypothesized two models: (i) the transcriptional activation occurred at 30% of the entire *A. thaliana* chromosomal regions; or (ii) stochastically at 30% of each insertion event. To test which model is suitable for this transcriptional activation, we analyzed the transcriptional behavior of *LUC* genes that were integrated into close neighboring locations (Fig 4A). Theoretically, *LUC* pairs inserted in close proximity could result in three transcriptional fates: expression of both *LUC* genes (Fate A); expression of one *LUC* gene and silencing of the other (Fate B); and silencing of both *LUC* genes (Fate C) (Fig 4B). If the transgene transcriptional activation depends on the chromosomal locus, the transcriptional fates of neighboring *LUC* inserts are expected to be similar (Fig 4C). Hence, in this scenario, only Fates A and C would be observed for the *LUC* pairs (Fig 4C). Moreover, the expected ratio between Fates A and C would be 30:70 (Fig 4C), assuming an average transcriptional activation rate of 30%. Conversely, as shown in Fig 4D, if the transgene transcriptional activation occurs stochastically at 30% of each integration event and is independent of the chromosomal locus, the distribution of the transcriptional fates of *LUC* pairs would fit the joint probability of two individual activation events. In this model, the distribution ratio among Fates A, B, and C would be 9, 42, and 49, respectively (Fig 4D). According to these expectations, we examined which activation model fits the transcriptional activation of promoterless *LUC* genes. In our dataset, we identified 21 genomic locations in which independent *LUC* inserts were integrated within a 50-bp sliding window. Among these 21 *LUC* insert pairs, all three possible transcriptional fates were observed, as follows: Fate A, three cases; Fate B, five cases; and Fate C, 13 cases (Fig 4E, upper panel). This distribution fits the integration-dependent stochastic transcriptional activation model (Fig 4D), rather than the chromosomal-locus-dependent model (Fig 4C). In fact, the expected values, i.e., Fate A (1.9 events), Fate B (8.8 events), and Fate C (10.2 events), were not significantly different from the observed rates (Fisher’s exact test, *P* = 0.55) (Fig 4F). To perform a more rigorous test of the stochastic transcriptional activation model, we reduced the sliding window to 10 bp, which yielded 12 genomic locations (Fig 4E, lower panel). Similar to the results of the 50-bp sliding window analysis, the pairwise *LUC* comparison did not detect significant differences (Fisher’s exact test, *P* = 0.82) between the observed and theoretical values (Fate A, 1.0 vs. 1.1; Fate B, 3.0 vs. 5.0; Fate C, 8.0 vs. 5.9) (Fig 4F). It should be emphasized that the individual *LUC* inserts used for this integration-site neighborhood analysis stemmed from different, independently transformed cells that passed through ~10 cell divisions before nucleic acid extraction. Thus, we concluded that the transgene transcriptional activation in a given genome location was likely to be the outcome of an integration-dependent stochastic phenomenon.

**Fig 4.**
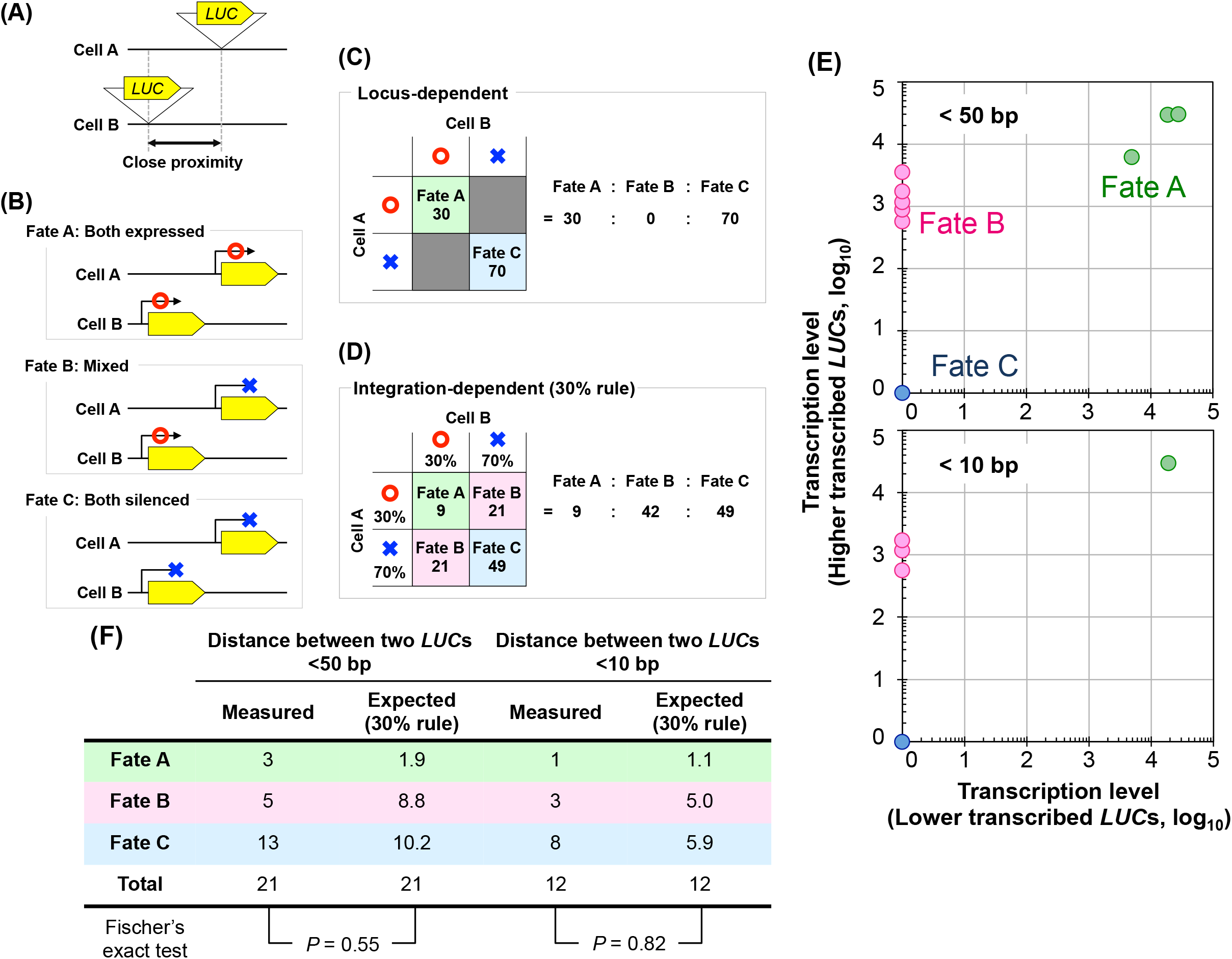
Transcriptional states of neighbouring *LUC* insert pairs located in close proximity. (A)*LUC* pairs inserted in close proximal chromosomal regions were used for integration-neighbourhood analysis. (B) Three possible fates of the transcription of *LUC* pairs: Fate A, expression of both *LUC* genes; Fate B, expression of one *LUC* gene and silencing of the other; and Fate C: silencing of both *LUC* genes. (C and D) Expected ratio of the three transcriptional fates classified in (B) for *LUC* pairs obeying (C) locus-dependent activation or (D) integration-dependent stochastic activation. (E) Transcriptional states of neighbouring *LUC* pairs inserted in the different cells. The distances between each neighbouring *LUC* insert were <50 bp (upper panel, n = 21) and <10 bp (lower panel, n = 12). (F) Measured and expected number of *LUC* pairs with Fate A, Fate B, and Fate C, as described in (E). The expected number was calculated according to the integration-dependent stochastic activation, assuming each transcriptional activation rate as 30%.

## Discussion

In this study, we performed a genome-wide screening of promoter-trapping events covering both expressed and unexpressed inserts for the first time, using a non-selective reporter. Collectively, the data revealed a new type of transgene transcriptional activation of the plant genome, which occurs stochastically at about 30% of each DNA integration event but not depending on the chromosomal loci. This transcriptional activation occurred in the transgenic cells that experienced only ~10 times cell divisions after the transgene integration, indicating that it is an immediate response of the plant genome to the incoming coding sequences. To date, we could not find any specific motifs that were enriched at the 5’ proximal regions of the transcribed *LUC* genes, which was quite a different situation from the annotated gene promoters (S10 Fig). How can we explain the mechanism of this new type of transcriptional activation that is stochastic and independent of the DNA sequences surrounding the transgene insertion sites?

It is generally accepted that T-DNA is integrated into the host genome following the double-stranded DNA breaks, which are repaired predominantly by the non-homologous DNA end-joining [22, 23]. This repair process remodels the chromatin and leaves so-called DNA damage scars in the chromatin epigenetic structure [24, 25]. This chromatin remodeling may account, at least in part, for the integration-dependent stochastic transcriptional activation. From this viewpoint, it is intriguing to compare the chromatin structures before and after the *LUC* integration, but this analysis remains technically challenging. In the present study, the established transgenic cell pools were highly heterogeneous; they contained thousands of distinct transgenic cell lines, and each cell line consisted of only ~1,000 cells. There is no practical methodology to analyze epigenetic configurations of each transgenic line from such a heterogeneous cell population. Recently, single-molecular resolution techniques were reported for the chromatin accessibility assay [26–28], but further technical breakthrough is needed to determine the epigenetic marks on the single chromatin molecule. In addition, the *LUC* mRNA level of each transgenic line was quite low compared with the annotated gene transcripts. Therefore, TSS determination of the *LUC* inserts is also challenging. We are now attempting these challenging analyses, which would provide useful information to depict the transcription initiation mechanism in this new type of plant genome response.

In the present study, we characterized a novel plant genome response to the incoming coding sequences, i.e., integration-dependent stochastic transcriptional activation. Contrary to the conventional gene-/promoter-trapping scenario in the HGT/EGT process, this foreign gene activation mechanism seems less harmful to the host nuclear gene networks because this mechanism does not cause disruption of the preexisting nuclear genes. Therefore, this finding provides a new angle for examining the gene activation mechanism in the massive gene transfer events between phylogenetically distant organisms. To evaluate the biological contribution of this novel genome response to plant genome evolution, further information is needed on how activated transcription via this mechanism continues and behaves over generations, and how selective pressure on the activated transcriptions affects their fates. Experimental studies along these lines could open the way to an understanding of how the initial molecular response of the eukaryotic genome is linked to the phenotypic evolution.

## Materials and Methods

### Construction of barcode-labelled plasmid libraries

The transformation vector plasmid was constructed using a modified pGreenII vector [29, 30] to encode 12 bp of random sequence (“barcode”), a promoterless firefly luciferase (*luc*+) coding sequence, a *nos* terminator sequence and an expression cassette of a kanamycin-resistant gene within the T-DNA region (S1A Fig and S1 Methods). All primers used in this study are listed in S1 Table.

### Plant cell culture and transformation

*A. thaliana* T87 cells [31] were cultured in mJPL3 medium [32] under continuous illumination (60 μE m^−1^ s^−1^) at 22°C with shaking (120 rpm). One-week-old cultures were collected using a 10 μm nylon mesh, washed with H_2_O twice and subjected to DNA, RNA and chromatin isolation and transformation.

*Agrobacterium tumefaciens* (GV3101) cells were transformed with the barcode-labelled libraries. *Agrobacterium*-mediated transformation of *A. thaliana* T87 cells was carried out according to the published protocol [32, 33]. We obtained three independently transformed pools of T87 cells (termed TRIP pools hereafter), which were grown on mJPL3 plates containing 25 μg ml^−1^ meropenem (MEPM) and 30 μg ml^−1^ kanamycin (Km) at 22°C under continuous illumination for about 2 weeks. Green calli were cultured in liquid mJPL3 medium containing 12.5 μg ml^−1^ MEPM and 10 μg ml^−1^ Km with shaking under continuous illumination at 22°C for 2 weeks. Finally, the cells were transferred to fresh mJPL3 medium and grown for 1 additional week.

### Determination of the insertion loci of LUC genes

Two micrograms of genomic DNA extracted from the TRIP pools using the DNeasy Plant Mini Kit (QIAGEN) were digested completely with *Dpn*II, purified using the QIAquick PCR purification kit (QIAGEN) and circularized with T4 DNA ligase. After purification using the QIAquick PCR purification kit, the circularized DNA was subjected to inverse PCR using primers that were designed to hybridize within the *LUC* gene (S1B Fig). From this point, we prepared two types of sequencing libraries: (1) The inverse PCR product was digested completely with *Apa*LI or *Sca*I to block the amplification of the vector-backbone-containing fragments in the subsequent steps. Nested PCR was performed, followed by sequencing library preparation using the Nextera XT DNA Sample Prep Kit (Illumina); (2) The inverse PCR product was subjected to tailed-PCR and digestion with *Apa*LI or *Sca*I, followed by the addition of terminal adapters via one additional round of PCR, to prepare the sequencing libraries essentially according to Akhtar *et al*. [16]. Sequencing was performed on an Illumina MiSeq sequencer with 301 bp paired-end reads.

The insertion loci of *LUC* genes were determined using an open-source software and custom Perl scripts (S1B, and C Fig). Briefly, the sequencing reads were trimmed from the 3’ end with a phred-scaled quality score ≥30. Reads containing a *LUC* segment (31 bp), the barcode (12 bp) and a *LUC* flanking sequence (25–50 bp) were extracted. The *LUC* flanking sequences were mapped to the TAIR10 version of the *A. thaliana* genome using Bowtie [34] with the following parameters; *bowtie −m 1 −v 3*. Subsequently, the 3’-junction sites of the mapped flanking sequences were defined as the genomic loci of the corresponding *LUC* inserts. Reliable locus–barcode pairs of *LUC* inserts were collected according to their read depth; at least three reads and 90% of individual mapped loci were occupied by an identical barcode sequence. We combined all *LUC* loci that were derived from three biologically independent TRIP pools, as well as from two of mapping libraries, and subjected them to subsequent analyses. For additional details, see S1 Methods.

### Determination of the relative transcription levels of *LUC* genes

RNA was extracted from the TRIP pool using the RNeasy Plant Mini Kit (QIAGEN) and treated with RNase-free DNase I (QIAGEN). cDNA was synthesized from 5 μg of the RNA using an oligo dT_15_ primer and SuperScript III Reverse Transcriptase (Thermo Fisher Scientific). Sequencing libraries were prepared by amplification of the barcode region using primers with an adapter extension, followed by tailed-PCR using Nextera XT Index Primers (Illumina) (S1B Fig). From an aliquot of DNA from the TRIP pools, sequencing libraries of the barcode region were prepared using the method described above. These cDNA and DNA libraries were sequenced on an Illumina MiSeq sequencer with 76 bp paired-end reads.

To determine the relative transcription levels of *LUC* genes, barcode sequences were extracted from sequencing reads and counted. Barcode sequences with a read number ≤5 in the DNA library were omitted. Moreover, barcode sequences with a read number ≤5 in the cDNA library were set as zero. For each library, the read number of each barcode was normalized to the total reads of the library. To obtain an indicator of the RNA level per DNA molecule, the cDNA read number was divided by the corresponding DNA read number and multiplied by 10,000, which was used to indicate the transcription levels of the individual *LUC* genes. For additional details, see S1 Methods.

### RNA-Seq

RNA was extracted using the RNeasy Plant Mini Kit (QIAGEN) and treated with RNase-free DNase I (QIAGEN). RNA-Seq libraries were prepared using the SureSelect Strand-Specific RNA-Seq Kit (Agilent), according to the manufacturer’s instructions. The libraries were sequenced on an Illumina MiSeq sequencer with 76 bp paired-end reads. The sequencing reads from two replicated experiments were combined. The transcribed regions and their expression levels were determined using STAR [35] and StringTie [36], with the *A. thaliana* genome (TAIR10) as a reference for mapping.

### ChIP-Seq

The fixation of *A. thaliana* T87 cells, chromatin isolation and fragmentation and ChIP (antibody: anti-H3K9me2 (MABI, 308-32361)) were performed basically as described by Saleh *et al.* [37]. Successful enrichment of ChIPed DNA was validated according to To *et al.* [38]. ChIP-Seq libraries were prepared using the DNA SMART ChIP-Seq Kit (Takara Clontech), according to the manufacturer’s instructions. Libraries were sequenced on an Illumina MiSeq sequencer with 76 bp paired-end reads. The sequences derived from a template-switching reaction were trimmed from the reads. Subsequently, the reads from two replicated experiments were combined and mapped to the *A. thaliana* genome (TAIR10) using Bowtie2 [39]. Peaks corresponding to H3K9me2 enrichment were called using MACS (version 2) [40].

## Acknowledgments

We thank A. Katahata, I. Ohshima and M. Mino for useful discussions. *Arabidopsis* T87 suspension-cultured cells (rpc00008) were provided by RIKEN BRC which is participating in the National Bio-Resource Project of the MEXT, Japan.

## Supporting information captions

**S1 Fig.**
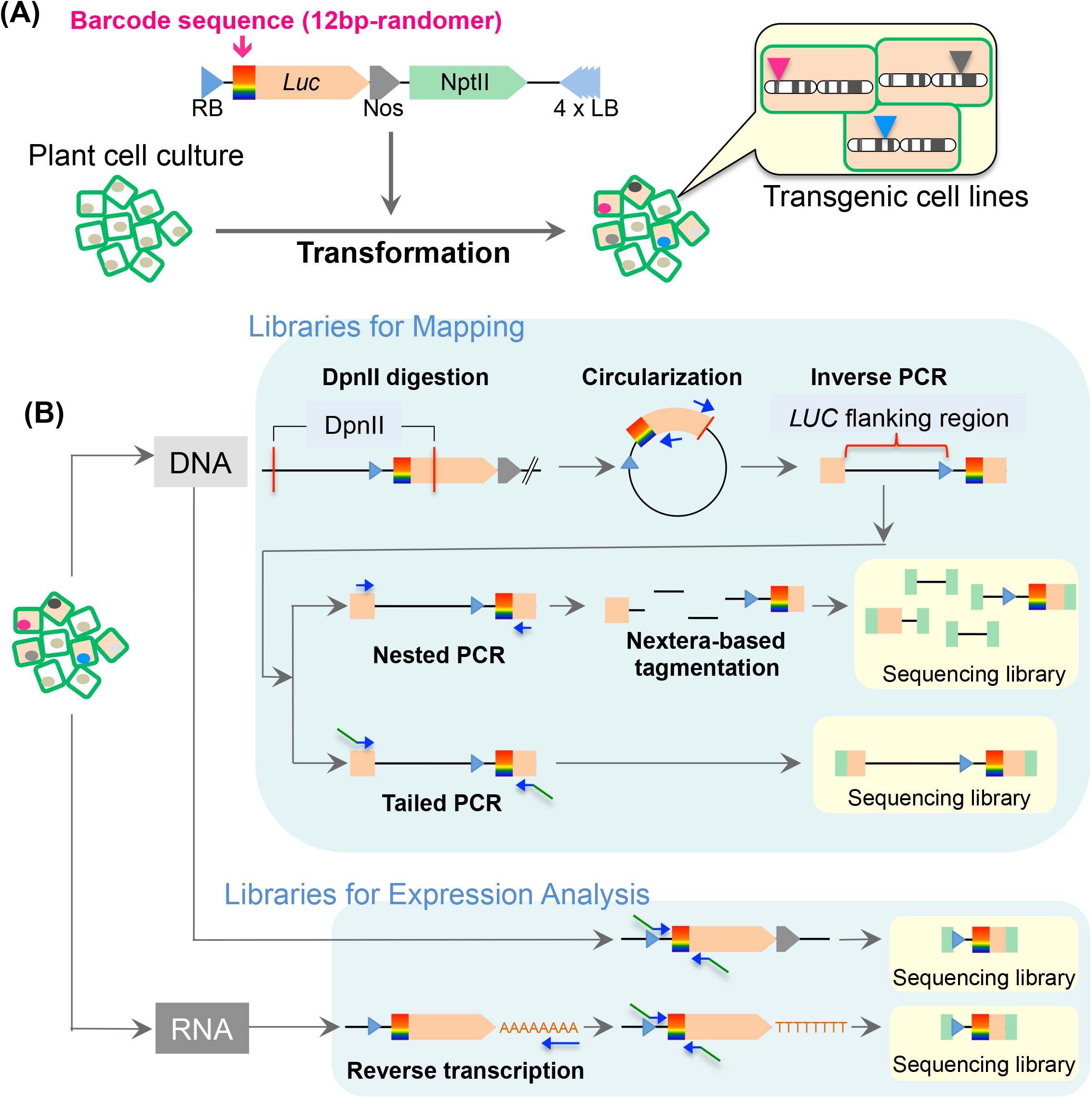
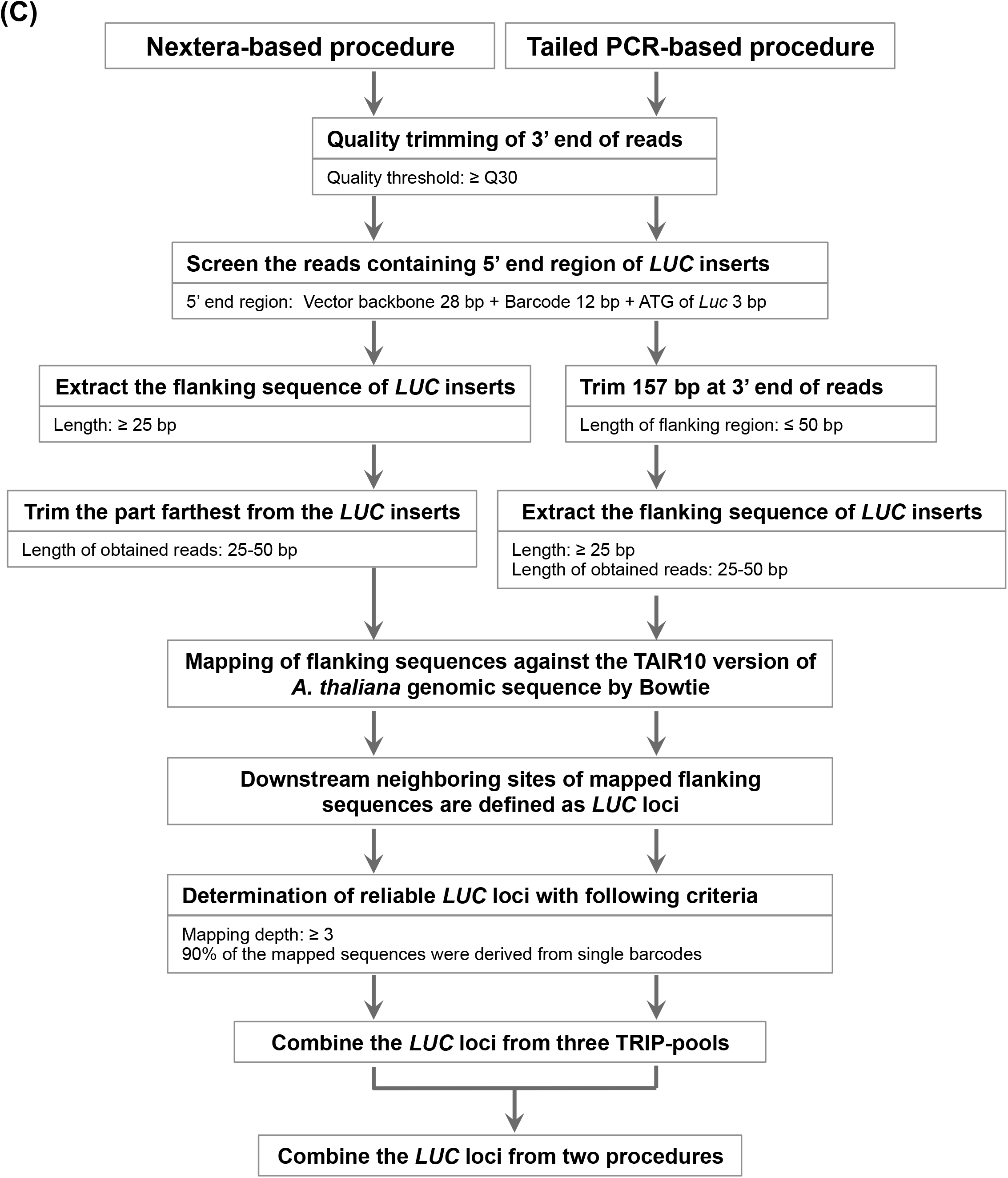
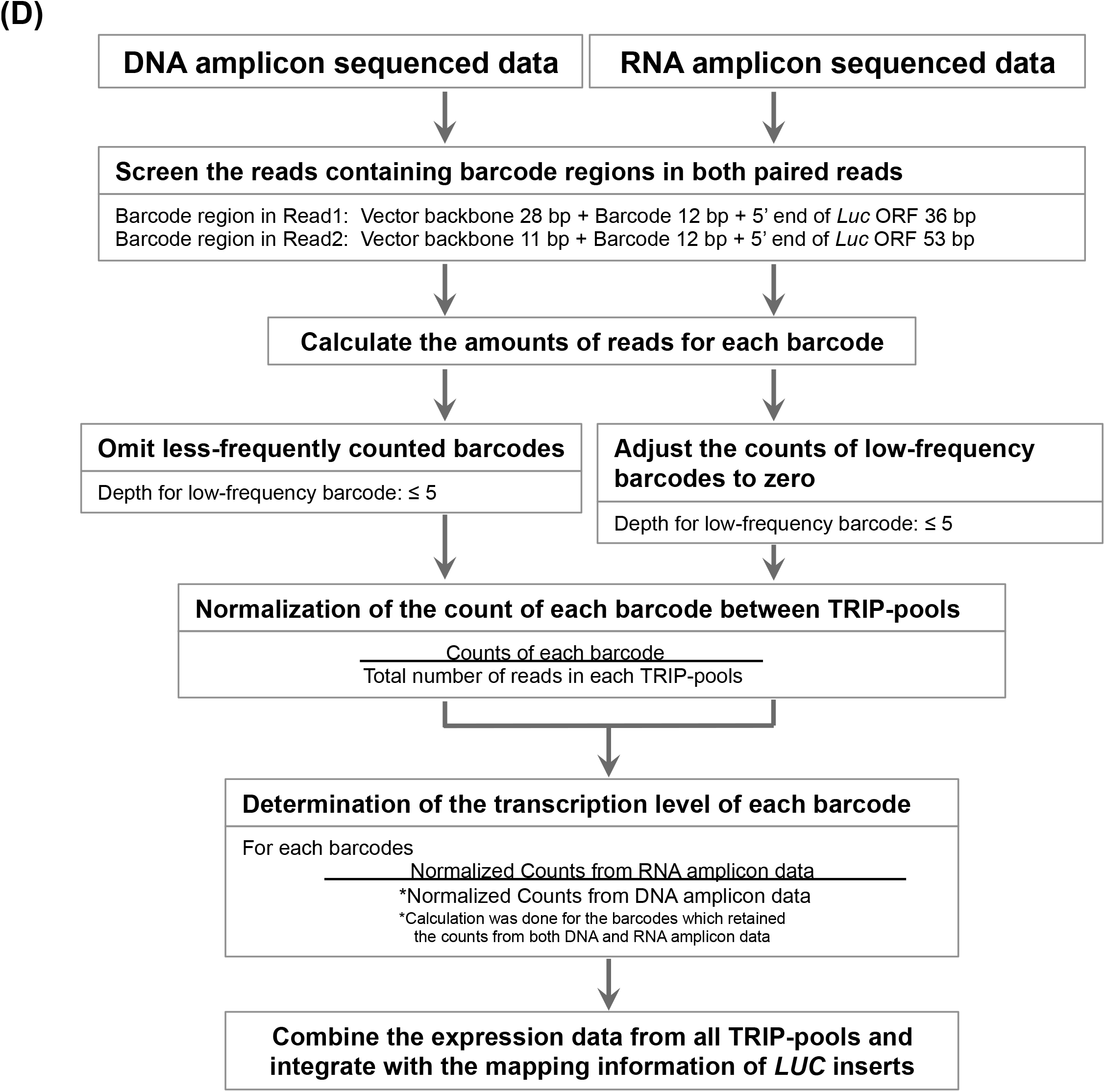
Precise workflow of the promoter analysis that was performed using the TRIP system. (A) Transformation of multiplexed barcoded vectors into *Arabidopsis* T87 suspension-cultured cells. (B) Preparation of sequencing libraries for the mapping and expression analyses. To prepare the mapping libraries, two different methods were employed after inverse PCR. In the first method, nested PCR products were fragmented and tagged with sequencing adapters using a Nextera-based method. In another method, inverse PCR products were subjected to tailed PCR, to add the sequencing adapters. To prepare libraries for the expression analysis, the barcode regions of both cellular DNA and cDNA were PCR amplified, followed by the addition of sequencing adapters using tailed PCR. cDNAs were prepared via an oligo(dT)-primed RT reaction. The libraries obtained were applied to a high-throughput sequencing analysis. (C) Workflow of the data-analysis pipeline that was used for the mapping of *LUC* genes. The flanking sequences of the *LUC* genes were extracted from the Nextera-based mapping library and tailed-PCR-based mapping library using slightly different methods. The *LUC* loci obtained were combined in the final step. (D) Flow diagram used for the determination of *LUC* transcription levels. The transcription level data obtained for individual barcodes were associated with the respective mapped *LUC* genes and used in subsequent analyses.

**S2 Fig.**
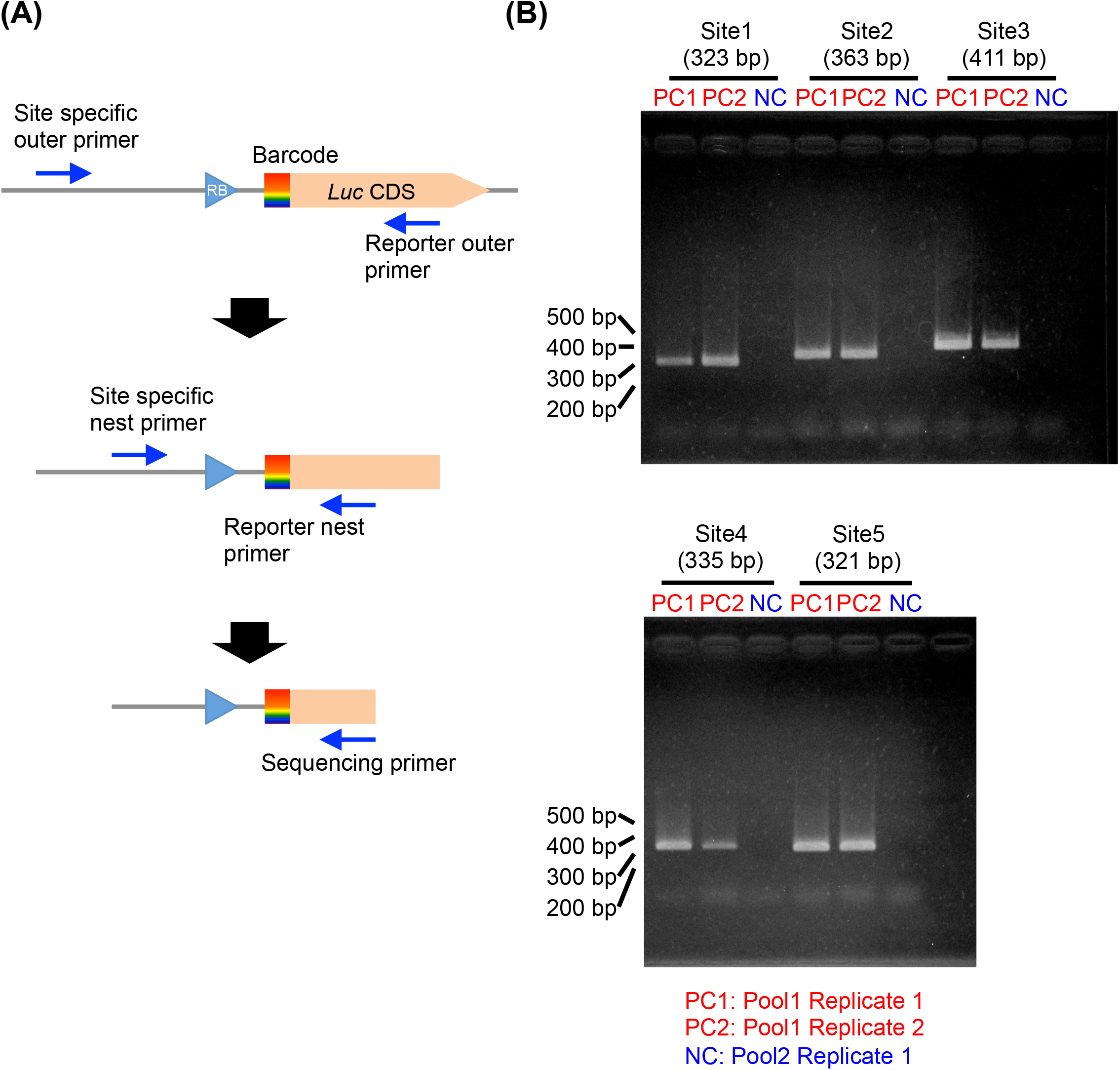
Validation of the *LUC* mapped loci and barcode sequences via PCR amplification in five representative samples. (A) Schematic diagram of the nested PCR that was performed using insertion-site-specific and *LUC*-specific primers. (B) Five *LUC* genes were chosen from the TRIP-Pool1 and detected by PCR. PC1 and PC2 are technical replicates of the PCR using the template DNA from TRIP-Pool1 cells. NC is the PCR product from the DNA of TRIP-Pool2 and was used as a negative control. The PCR products were loaded onto a 2% agarose gel. The expected size of the PCR products is shown at the top of the gel, in parentheses. The PCR products obtained were Sanger sequenced for verification of the barcode sequences.

**S3 Fig.**
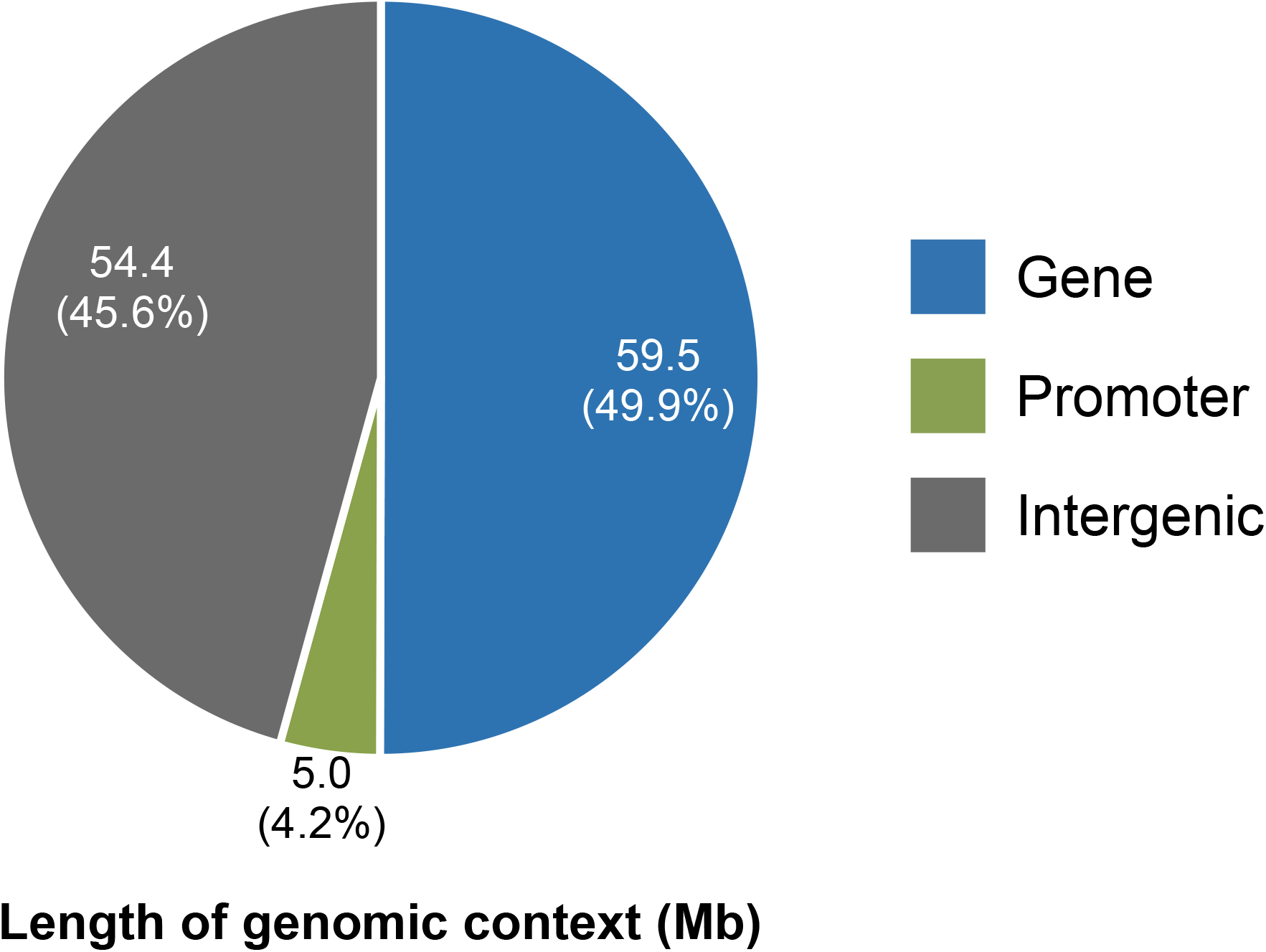
Length of each genomic context. The total length of the respective genomic contexts and their percentage in the whole genome are shown. The 200 bp segments 5’-proximal to the genic region (CDS plus UTR regions according to TAIR10) were defined as promoter regions, and the remaining sequences were defined as intergenic regions. When neighboring promoter and genic regions were overlapped, those parts were omitted from the statistical analyses described above (their sum was 0.23 Mb, 0.2% of the whole genome).

**S4 Fig.**
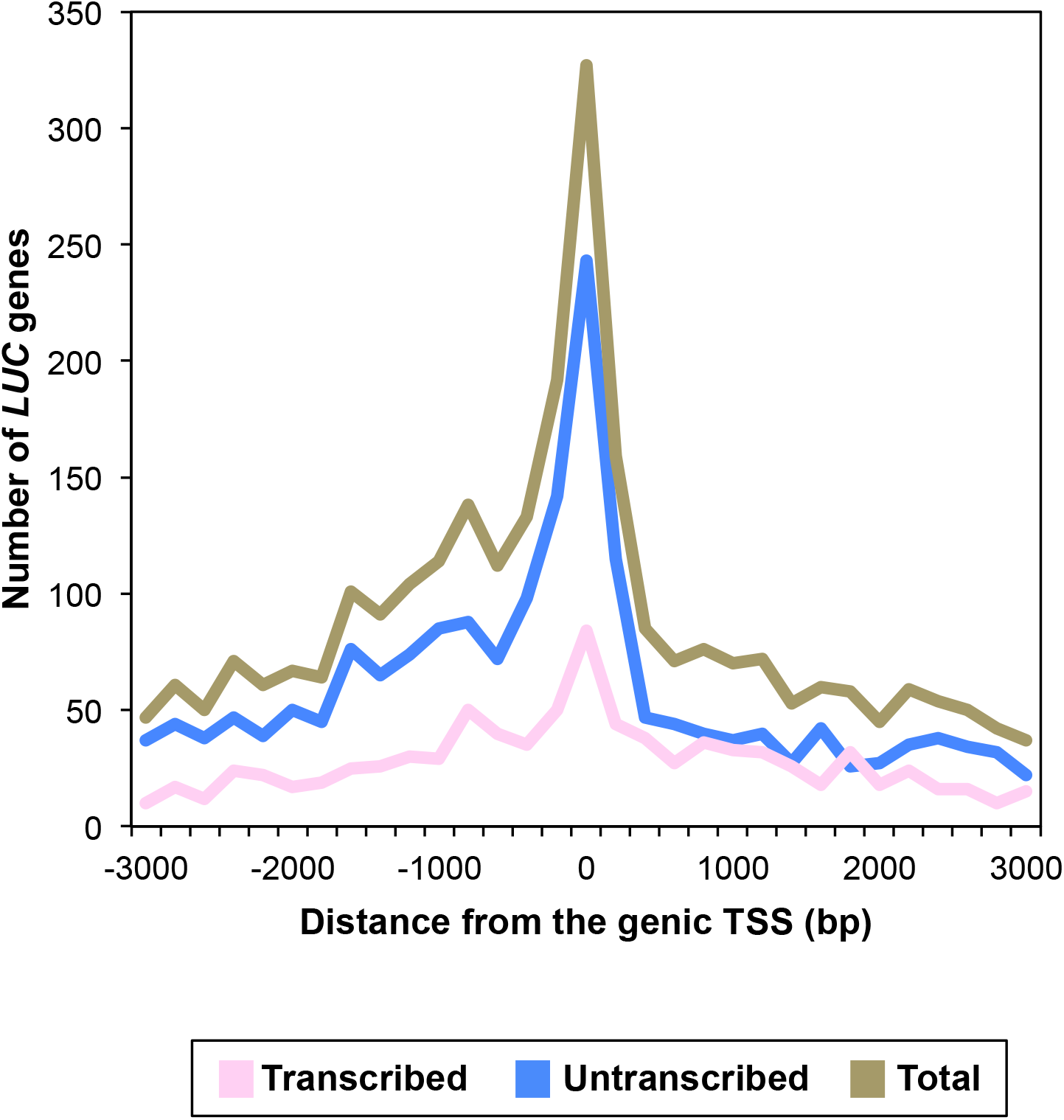
Abundances of *LUC* genes relative to the nearest genic TSS. Number of LUC genes in relation to the distances from the genic TSS was counted in 200 bp window size.

**S5 Fig.**
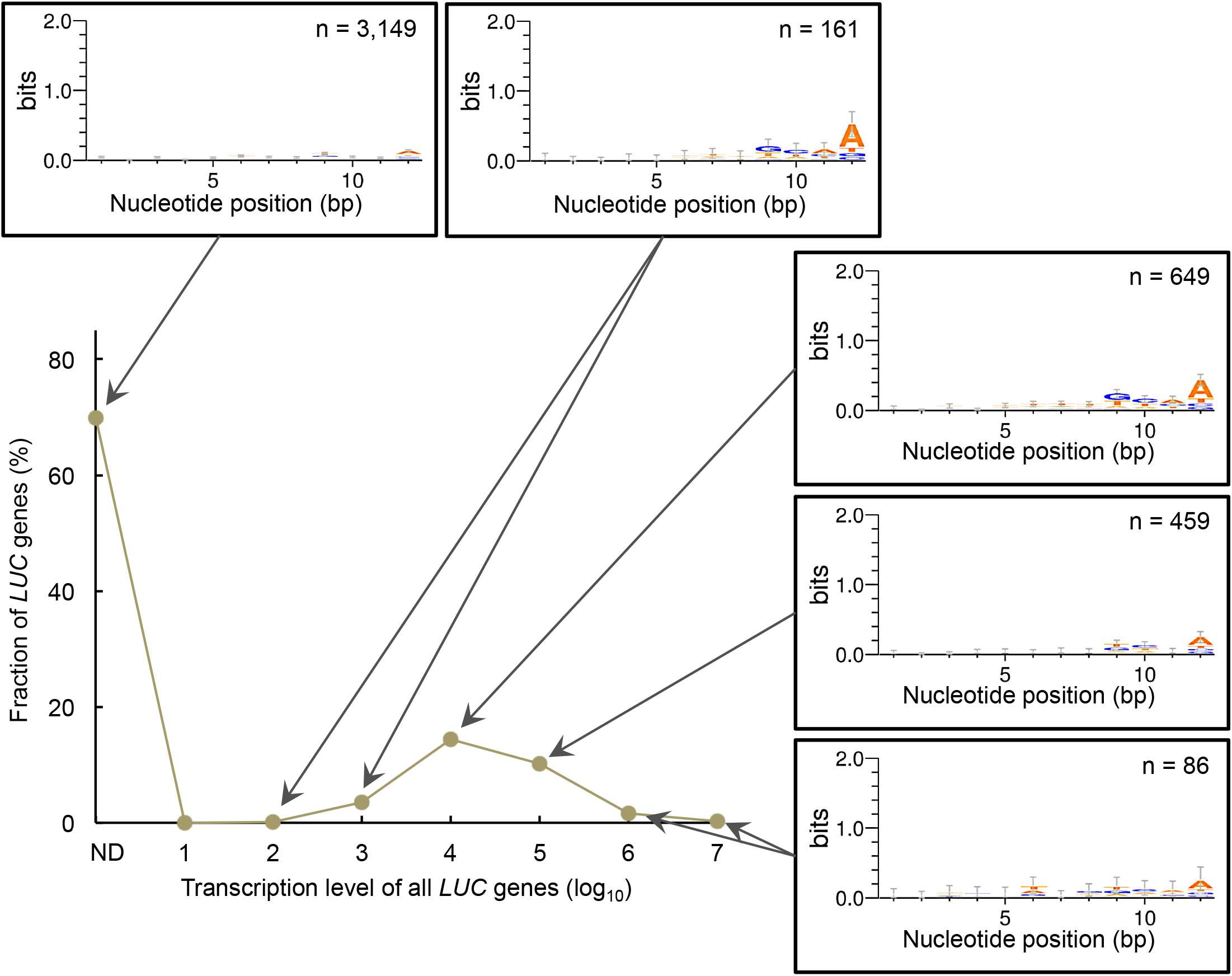
Assessment of the effect of barcode sequences on the LUC transcription levels. Frequently observed barcode motifs in the LUC insert of indicated transcription levels were analyzed using WebLogo3 (Crooks *et al.*, 2004). The transcription levels of all the LUC genes are shown as in Fig 1D. A weak positional preference for ‘A’ was found at the 3’-terminal position on the barcode. However, the frequency of ‘A’ at this position did not correlate with the strength of transcription.

**S6 Fig.**
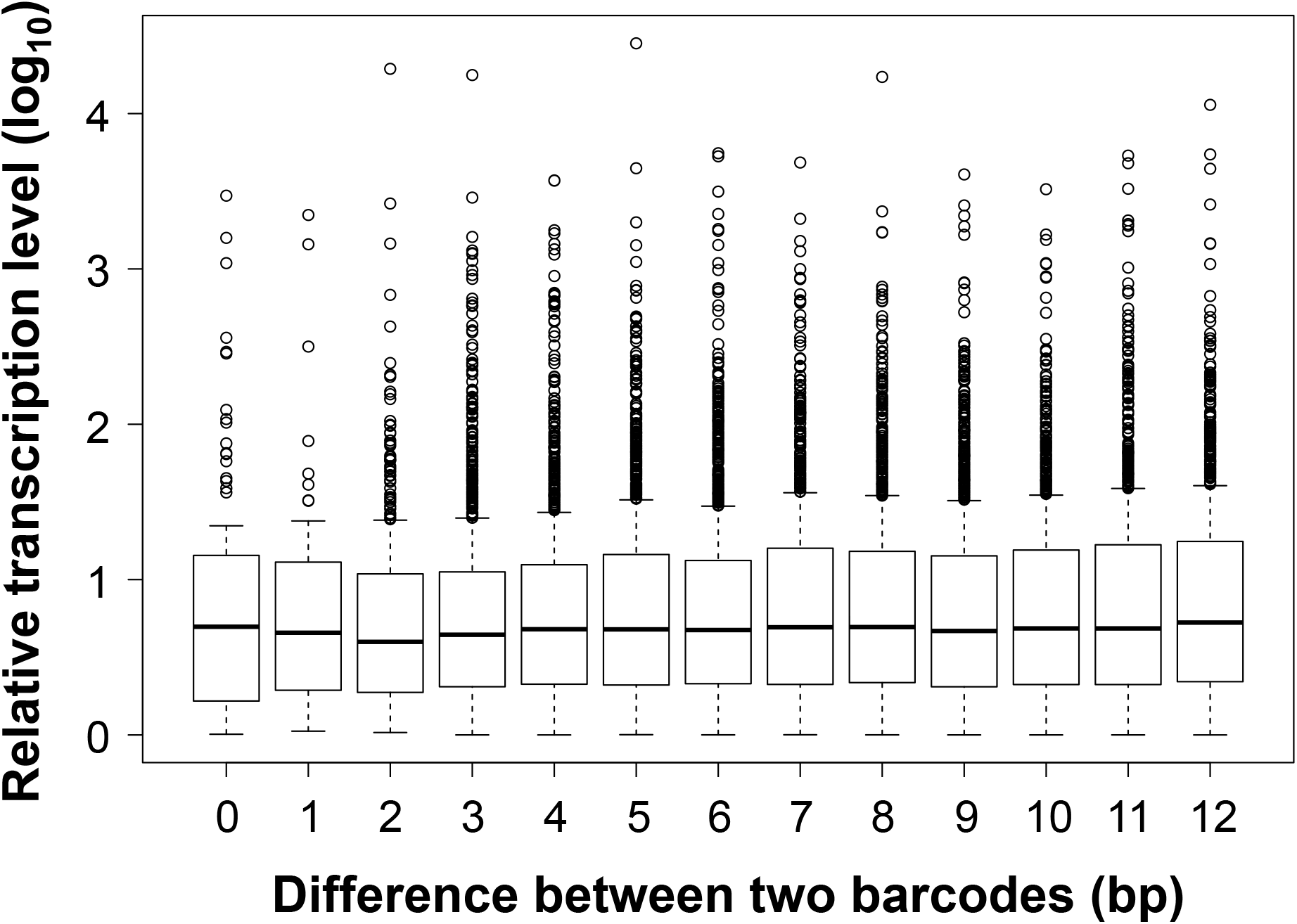
Similarity/dissimilarity of the transcription levels of the randomly selected LUC pairs against the sequence identity of the 12-base barcode. A pair of LUC genes was randomly selected from the 4,504 mapped LUC genes, and the similarity/dissimilarity of their transcription levels is shown as the ratio of their RNA levels in a logarithmic scale; the ratio was calculated by dividing the higher RNA level by the lower level (i.e., log(ratio) ≥0). The similarity/diversity of the barcode is indicated by the number of mismatched nucleotides at the corresponding positions. This graph is the summary of the analysis of 10,566 LUC pairs and indicates the absence of a correlation between the similarity of the barcode sequence and that of the transcription level. In other words, the barcode sequence does not affect the transcription level of LUC genes. Methods note: 1) When randomly selected LUC pairs were located within 100 kb on the same chromosome, they were omitted from the analysis, lest their positional effect should influence their transcription levels. 2) One thousand LUC pairs were analyzed each for the indicated number of mismatches in the barcode. However, for mismatch numbers of 0, 1, and 2, the number of LUC pairs analyzed was 92, 51, and 423, respectively. This is because the number of such highly homologous barcodes in the total population of 4,504 LUC inserts was limited, and these are all the LUC genes that fulfilled the given requests. 3) The LUC inserts of the identical barcodes were derived from different TRIP pools, because LUC mapping in a given TRIP pool had been conducted so that the individual LUC genes were mapped to a unique locus, with omission of those that were mapped to more than one locus.

**S7 Fig.**
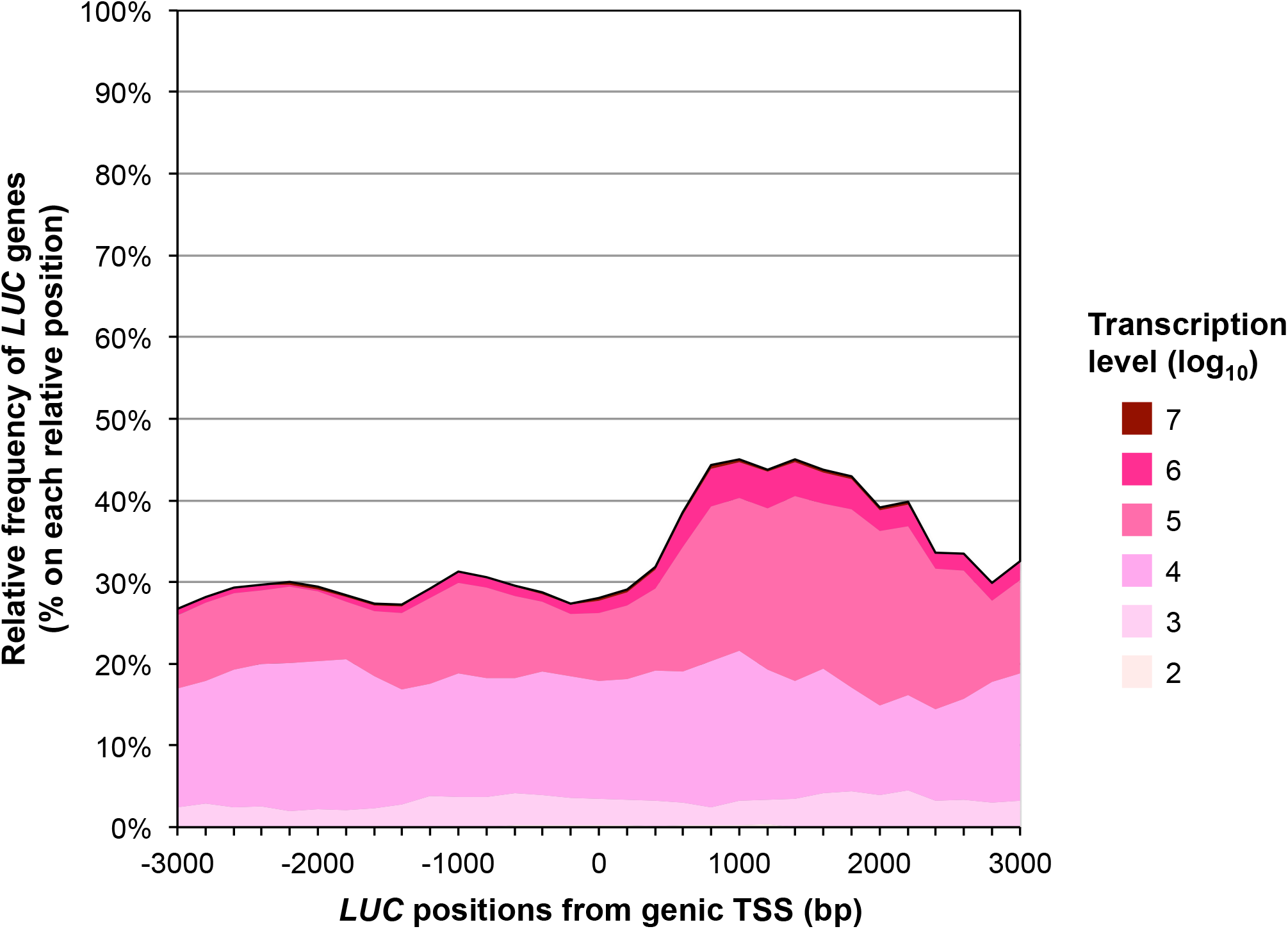
Frequency of transcribed *LUC* genes relative to the annotated genic TSS. Abundance of the *LUC* genes with the indicated transcription levels in relation to the distance from the genic TSS, as shown in Fig 1E. The plot was smoothed by calculating the five-point moving average of integration frequency in each window (200 bp).

**S8 Fig.**
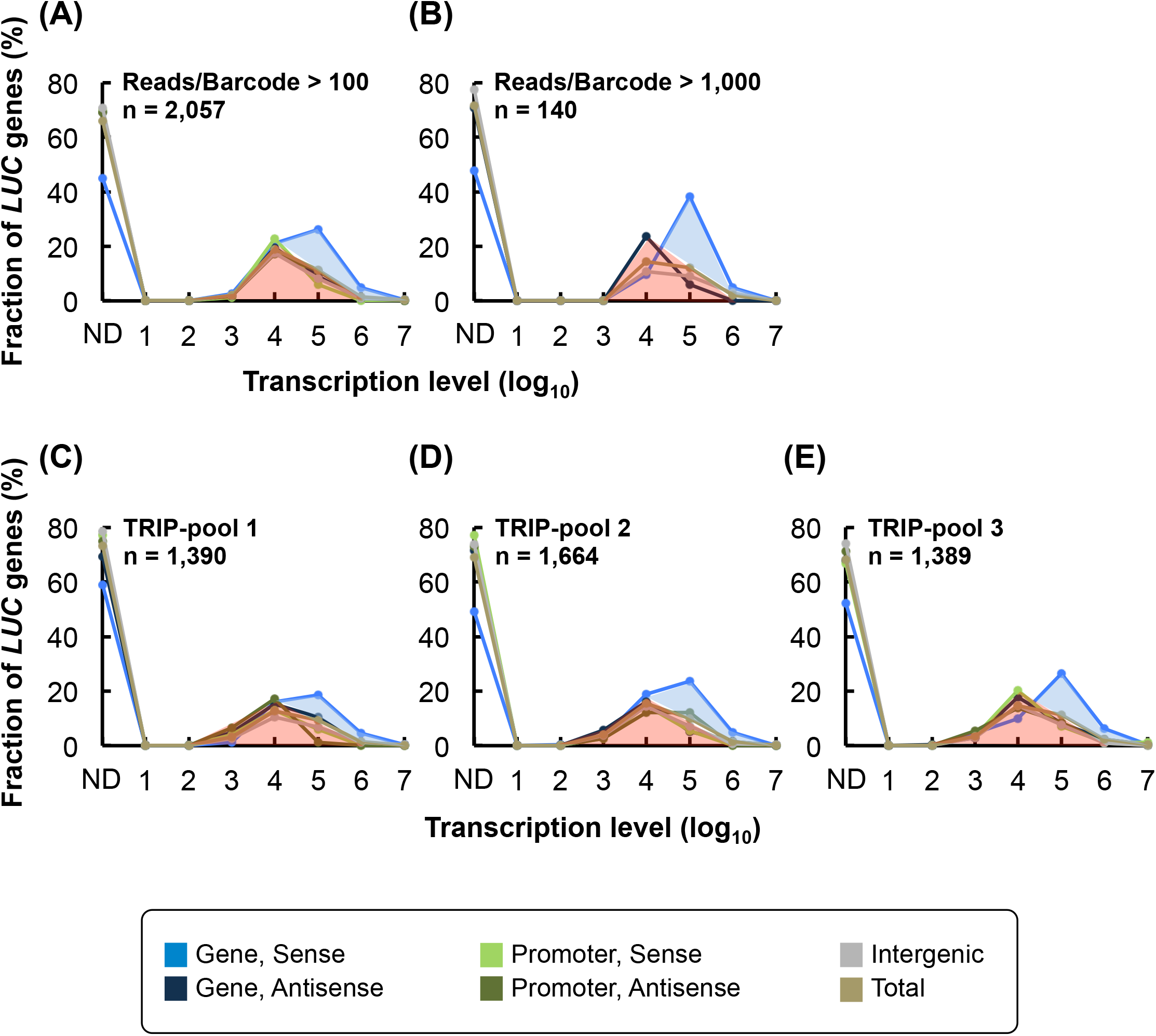
Expression profiles of *LUC* genes with high-number reads from the amplicon-sequencing data and of *LUC* genes from biological replicates. (A and B) For each barcode, when the number of reads from DNA amplicon sequencing was up to (A) 100 or (B) 1,000, the barcode was omitted from the analysis. The number of reads for each barcode obtained from RNA amplicon sequencing was redefined as zero, if the number of reads was below such thresholds. The subsequent processes used in this analysis were same as those used in Fig 1D. The expression profiles of the *LUC* genes located in promoter regions were omitted from (B), because the number of such *LUC* genes was insufficient to represent their profiles. (C–E) Expression profiles of three biological replicates. The numbers of *LUC* genes shown in all graphs are the total amount of *LUC* genes used for their analysis. The fraction of the transcribed *LUC* genes attributed by two distinct mechanisms are indicated by light-blue and light-red areas.

**S9 Fig.**
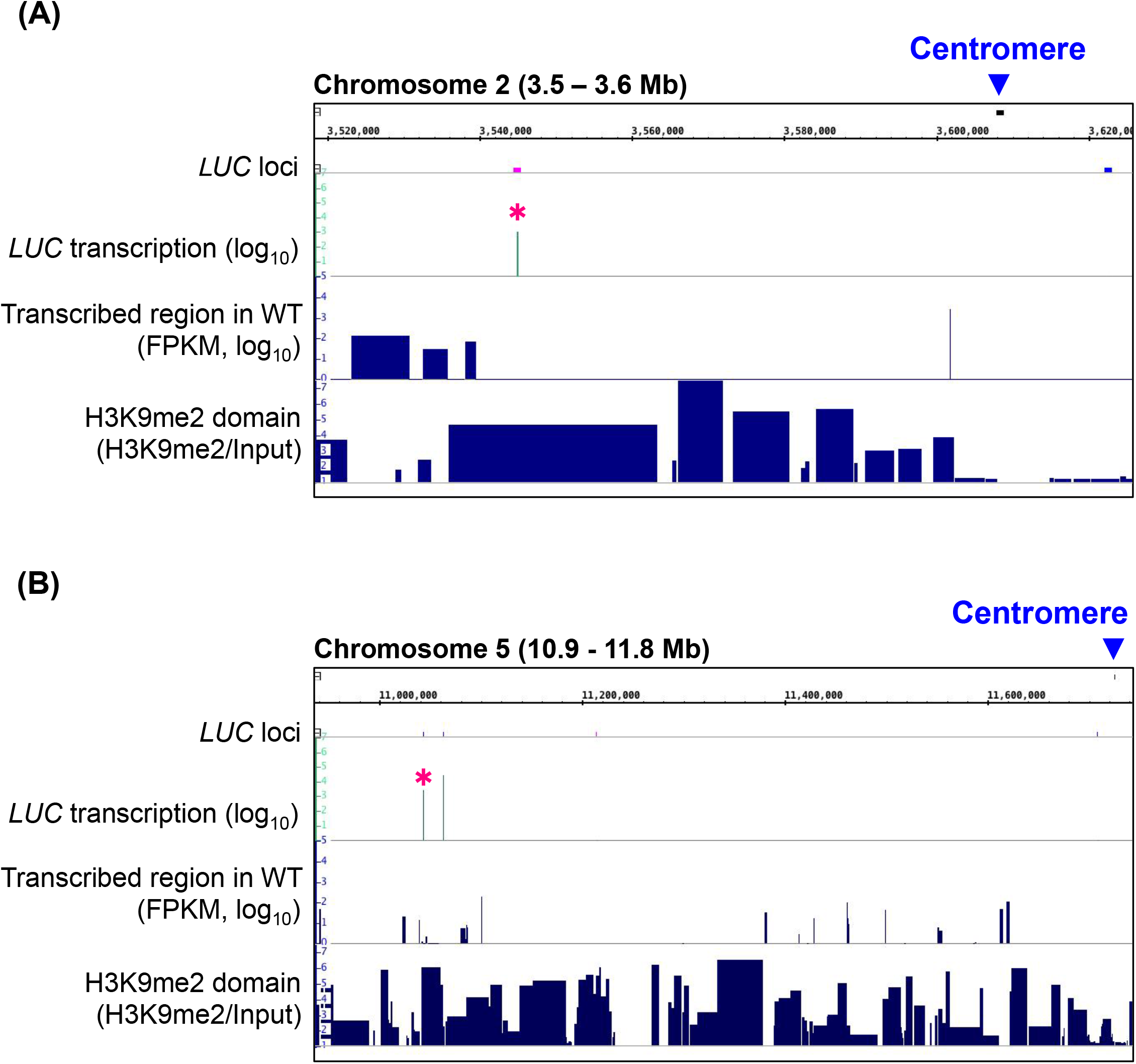
Two examples of transcribed LUC genes in the H3K9me2-marked regions located around the centromere. (A and B) Transcribed LUC genes (asterisk) were found 63 kb and 682 kb away from the centromeres of chromosomes 2 (A) and 5 (B), respectively. The respective H3K9me2 levels of these loci were 80 (A) and 91 (B) percentiles, respectively. In WT T87 cells, transcripts were very scarce in these heterochromatic regions.

**S10 Fig.**
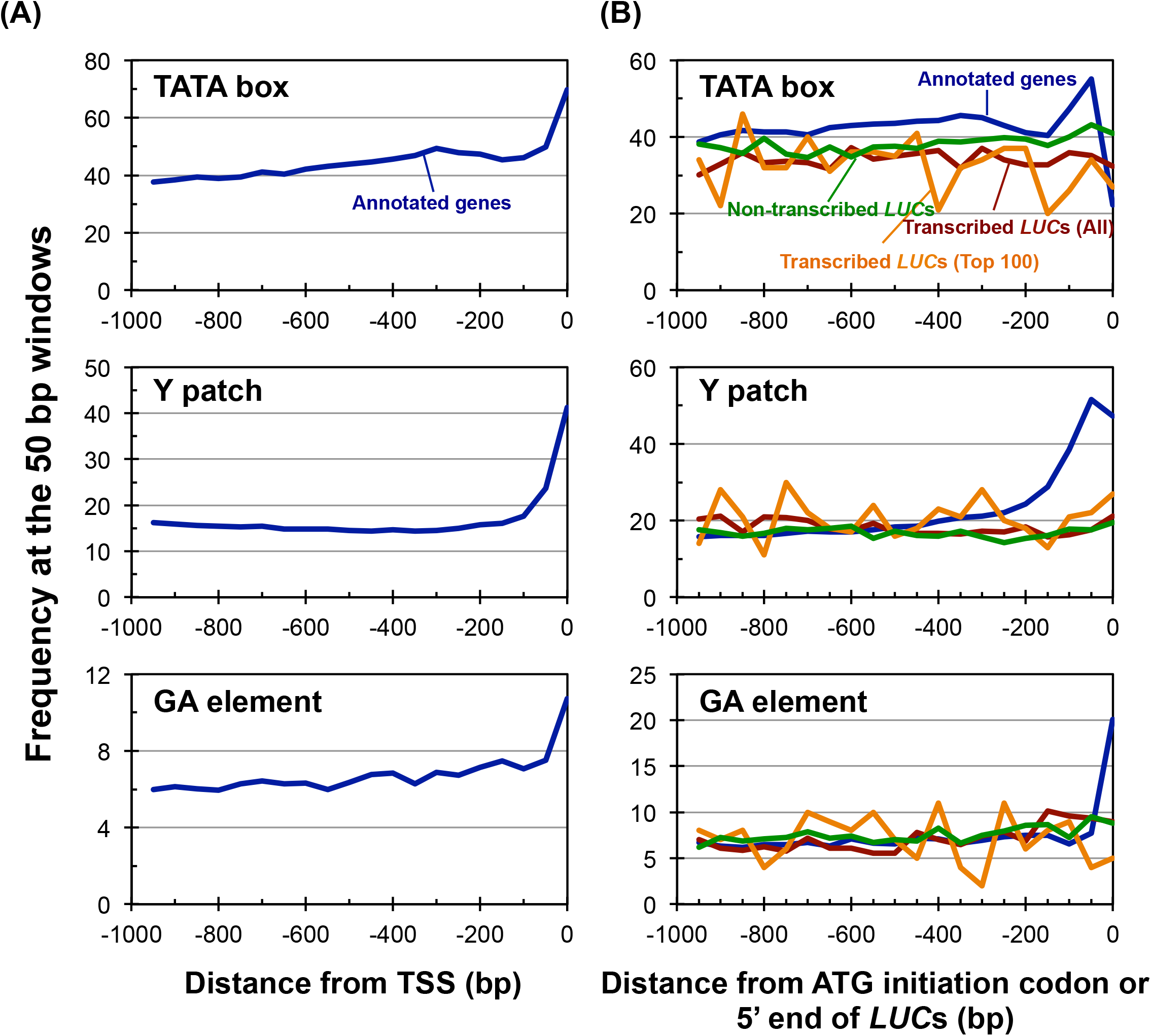
Distribution of *cis*-regulatory elements in the upstream region of *LUC* integration sites. (A and B) The frequency of TATA-box, Y patch, and GA elements in the upstream region of the (A) TSS of annotated genes, or (B) of ATG initiation codons of annotated genes and 5’ ends of *LUC* inserts were analyzed according to Yamamoto *et al.* (Yamamoto *et al.*, 2007) using a window size of 50 bp for the high-sensitive detection of the motifs. The Y-axis represents the fraction of genes or *LUC* genes that contained the indicated motifs.

**S1 Table.**
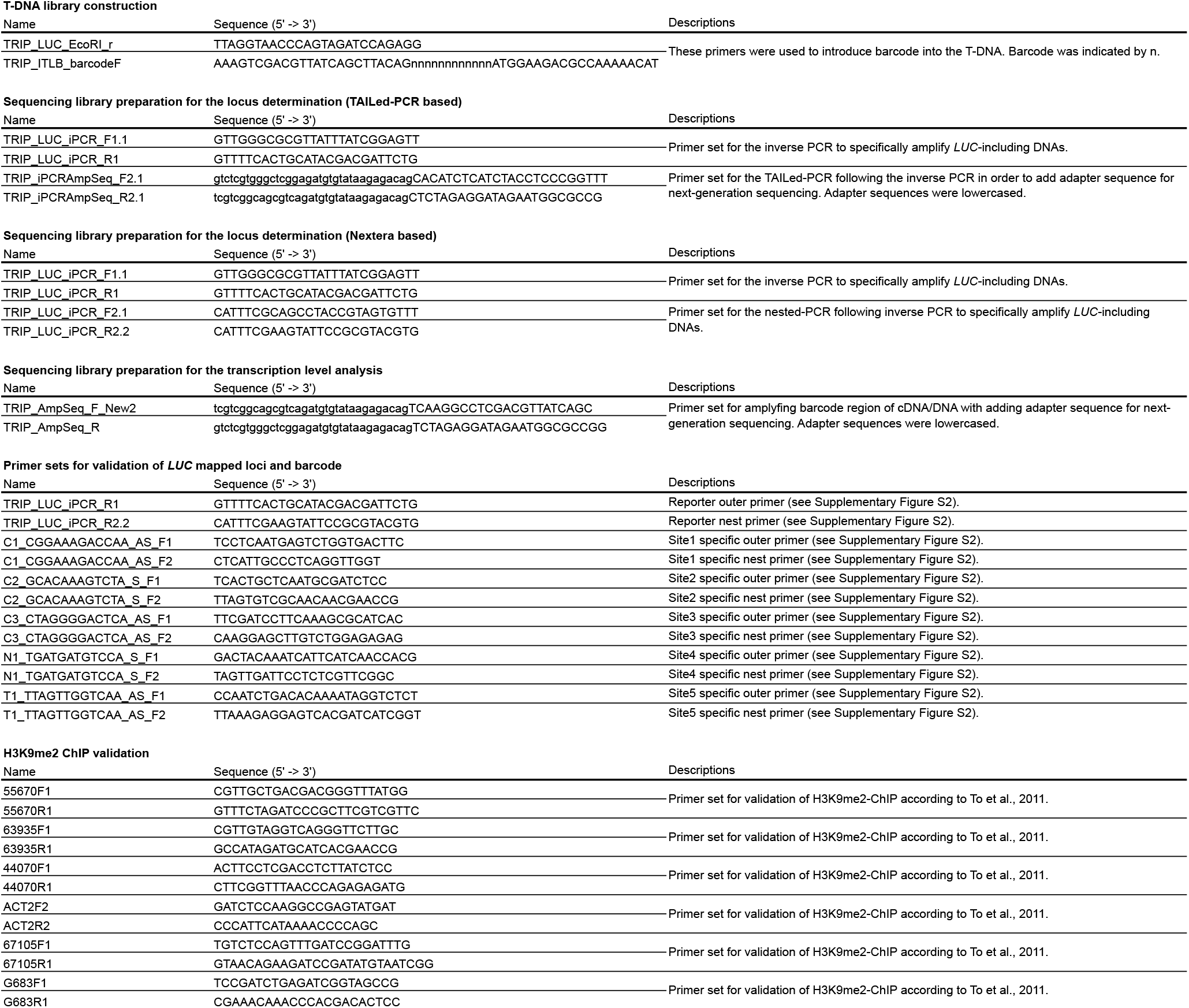
Primer list

**S1 Methods. Detailed methods for the massive promoter-trapping experiment**

## S1 Methods

### Construction of barcoded plasmid libraries

The transformation vector plasmid was constructed using a modified pGreenII vector [1, 2] to encode a promoter-less reporter cassette and a kanamycin-resistant cassette between the right (RB) and left border (LB) (S1A Fig). The reporter cassette consisted of a 12-base random barcode sequence, the firefly luciferase (*LUC*^+^) gene, and a *nos* terminator sequence, and contained the short sequence 5’–AGGCCTCGAGGTTATCAGCTTACAG–3’ (the *Xho*I site is underlined) between the RB and the random barcode. This short sequence was inserted for the sake of introducing the barcode sequence and also for the construction of amplicon-sequencing libraries. The kanamycin-resistant cassette contained the *NptII* gene with a *nos* promoter and *nos* terminator. The LB was modified to be repeated four times (S1A Fig), to suppress the integration of the vector backbone sequence into the plant genome [3]. The modified LB sequence was 5’–ATCCTGCCAGTTACACCACAATATATCCTGCCAGTTACACCACAATATATCCTGCCAGTTACACCACAATATATCCTGCCAGTTACACCACAATATATCCTGCCA–3’, and the first 9 bases were added through the construction step. To obtain a plasmid library that contained the random barcode sequence, the 5’-end fragment of the luciferase gene was amplified using two primers (5’–AAA<u>GTCGAC</u>GTTATCAGCTTACAGNNNNNNNNNNNNATGGAAGACGCCAAAAACAT–3’′and 5’–TTAGGTAACCCAGTAGATCCAGAGG–3’ (the *Sal*I site is underlined)), digested with *Sal*I and *Eco*RI (the *Eco*RI site was located on the amplified *LUC* fragment), and inserted into the *Xho*I and *Eco*RI sites of the transformation vector.

The constructed vector was transformed into *Escherichia coli* strain NEB 10-beta (New England Bilabs) by electroporation, and approximately 420,000 transformant colonies were obtained; this number suggests the initial diversity of the barcode clones. The transformed *E. coli* cells were cultured in liquid LB medium and subjected to plasmid DNA extraction.

### Mapping of the *LUC* genomic loci

The high-throughput sequencing data of the mapping libraries were trimmed from the 3’ end using fastq_quality_trimmer (http://hannonlab.cshl.edu/fastx_toolkit/) with a phred-scaled quality score ≥30 and were used for the mapping of *LUC* genes with the aid of open-source software and custom Perl scripts (S1C Fig). In *Agrobacterium*-mediated DNA integration, the 3’-terminal 3 bp of the RB is usually the junction between the T-DNA and the plant genome [4]. Therefore, the transformation vector sequence from the 3’-terminal 3 bp of the RB to the ATG initiation codon of *LUC* was used as the *LUC* segment, to search for the *LUC* flanking genomic sequences. The searching methods were slightly different between the two types of mapping libraries. (i) In the Nextera-based libraries, both paired-end reads were used to obtain *LUC* flanking sequences. Sequenced reads that included the *LUC* segment plus more than 25 bp of its flanking sequence were screened, and the flanking sequences and their corresponding *LUC* barcodes were extracted. The flanking sequences obtained were trimmed up to 50 bp using fastx_trimmer (http://hannonlab.cshl.edu/fastx_toolkit/). (ii) In the tailed-PCR libraries, only forward reads from the paired-end reads were used for the extraction of *LUC* flanking sequences, and their 3’-terminal 157 bp segments were removed using fastx_trimmer. From the obtained reads, 25-bp flanking sequences of the *LUC* segments and their corresponding barcodes were extracted as described above.

The flanking sequences obtained above were mapped to the *Arabidopsis* genome of TAIR10 version using Bowtie [5], on the condition that, at most, three mismatches were allowed and that individual sequences were associated with a unique genomic locus (Bowtie settings, −m 1, −v 3). Subsequently, the 3’-junction sites of the mapped flanking sequences were defined as the genomic loci of the corresponding *LUC* insertion sites. We also applied the following rules: 1) *LUC* genes that mapped at a single locus but for which the sequence reads were less than 3 were discarded and 2) cases in which very similar barcodes were mapped to identical loci (data not shown) suggested that an error occurred in the high-throughput sequencing of the barcode. Therefore, barcodes that occupied more than 90% of the reads at their respective genomic loci were retained, and their *LUC* genes were mapped to the respective loci.

Finally, we combined all *LUC* loci that were derived from three biologically independent TRIP pools, as well as from two kinds of mapping libraries, and subjected them to the following analyses.

### Determination of the transcription levels of *LUC* genes

Bioinformatics analyses of the *LUC* expression data were performed using a custom Perl script, the Microsoft Excel software, and the R package (http://www.R-project.org).

To compare the expression levels of individual *LUC* genes, their relative transcript levels were determined as follows (S1B and D Fig). The cDNA and DNA libraries that were specifically prepared for the expression analysis were subjected to amplicon sequencing on an Illumina MiSeq sequencer with 76 bp pair-end reads. The barcode sequences obtained were verified by the corresponding reads of each pair-end. The read numbers of each barcode sequences was counted in each sequencing library. We should note that, after sequencing on the MiSeq apparatus, *LUC* genes with a DNA read number ≤5 were omitted from the subsequent analysis. Besides, when the cDNA read number was ≤5, the transcript levels of the *LUC* genes were set as zero. Subsequently, the cDNA and DNA read numbers of the individual barcodes were normalized to the total cDNA and DNA read numbers of all barcodes, respectively. Then, the normalized cDNA barcode number was divided by the corresponding normalized DNA barcode number, to give an indicator of the RNA level per DNA molecule. This indicative number was multiplied by 10,000 and was used to indicate the transcription levels of the individual *LUC* genes. Obtained transcription levels of each *LUC* gene were then assigned to individual insertion loci described above according to the barcode sequences. *LUC* loci were omitted from subsequent analysis when transcription levels were not assigned.

## Notes

### Competing Interest Statement

The authors have declared no competing interest.

### Summary of Updates

Section on Abstract, Introduction, Results, and Discussion updated to clarify; Figures 1 and 2 revised; Supplemental Figures revised.

